# Molecular glue degraders of CK1α remodel lysosomes and limit ferroptosis in glioblastoma stem-like cells

**DOI:** 10.64898/2026.10.23.760392

**Authors:** Jane Jardine, Margaux Le Guyon, Kilian Trillet, Audrey Lacquemant, Isabelle Verdu, Talha Sagheer, Laura Merlet, Rosalie Moreau, Gwennan André- Grégoire, Mathilde Kerhervé, François Guillonneau, Guillaume Bastiat, Julie Gavard, Nicolas Bidère

## Abstract

Glioblastoma stem-like cells (GSCs) are highly plastic tumor cells that drive initiation, progression and therapeutic resistance. Although lysosomal homeostasis has emerged as a critical determinant of GSC fitness, the molecular mechanisms regulating this compartment remain incompletely understood. Here, using complementary proteomic approaches, we identified a lysosomal signature associated with the kinase CK1α in THP1 monocytic cells. Biochemical analyses revealed that genetic or siRNA-mediated depletion of CK1α altered the overall abundance and maturation of late endosome-lysosome-associated proteins in both THP1 cells and two patient-derived GSCs. This lysosomal remodeling was associated with impaired tumorsphere formation capacity in GSCs. To further explore the therapeutic potential of targeting CK1α, we employed selective molecular glue degraders (MGDs). Acute CK1α degradation efficiently recapitulated lysosomal remodeling and impaired tumorsphere formation in GSCs. While CK1α-targeted MGDs did not sensitize GSCs to lysosomal membrane permeabilization induced by L-leucyl-L-leucine methyl ester (LLOMe), they protected from GPX4 inhibitor-induced ferroptosis. Together, these findings identify CK1α as a regulator of lysosomal fitness and ferroptotic sensitivity, suggesting a connection between CK1α-dependent lysosome remodeling and associated cell death pathways.

## Introduction

Glioblastoma is the most common and aggressive brain tumor in adults. Despite surgical resection followed by radiotherapy and temozolomide chemotherapy, median survival remains approximately 15 months, and virtually all tumors recur [1, 2]. A major hypothesis for the recurrence rate is the presence of glioblastoma stem-like cells (GSCs), a highly plastic tumor cell population capable of sustained self-renewal [3]. These cells enable tumor initiation, growth and propagation [4–6], resist conventional radio/chemotherapies, and therefore may ultimately contribute to disease recurrence [7–9]. Therefore, identifying molecular mechanisms to increase GSC vulnerability is a critical challenge for developing more effective therapeutic strategies.

In recent years, lysosomes have emerged as an Achilles’ heel for GSC survival. Beyond their canonical catabolic functions, lysosomes serve as hubs for signaling, metabolism, membrane trafficking, and cell fate decisions [10]. One of the pioneer studies in this field reported that lysosomal remodeling through the action of the RNA- binding protein Quaking (QKI) could control the niche-independent self-renewal of GSCs [11]. Lysosomal lipid metabolism is also essential for GSC survival, as disruption of polyunsaturated fatty acid trafficking or cholesterol homeostasis through the inhibition of fatty-acid binding protein FABP3 or mucosa-associated lymphoid tissue lymphoma translocation-1 paracaspase (MALT1), respectively, compromises lysosomal membrane integrity and promotes lysosomal membrane permeabilization (LMP)-mediated cell death [12–14]. Together, these findings highlight the importance of lysosomal homeostasis for GSC maintenance and suggest that lysosomal vulnerabilities could be exploited therapeutically.

Ferroptosis is an iron-dependent form of cell death characterized by the peroxidation of membrane lipids [15]. It has attracted considerable interest as a strategy to target therapy-resistant cancer persister cells [16]. Increasing evidence points to an important role for lysosomes in ferroptosis, with inhibition of lysosomal acidification reported to suppress ferroptotic cell death [17, 18]. Consistent with this, lipid peroxidation initiates primarily at lysosomes during ferroptosis triggered by the inhibition of the antioxidant GPX4 [18]. This is in line with previous studies showing that manipulating lysosomal iron sequestration or activation can promote lipid peroxidation and ferroptotic death [19, 20]. More recently, the development of Fentomycin-1, a lysosome-selective iron-dependent phospholipid degrader, demonstrated that selective targeting of lysosomal iron was sufficient to trigger potent ferroptosis, further supporting a central role for lysosomal iron in ferroptotic cell death [18].

Casein Kinase 1 alpha (CK1α) is a pleiotropic kinase, with well-established roles in the regulation of β-catenin signaling, p53, or antigen receptor signal transduction [21–25]. Increasing evidence also implicates CK1α in lysosome-related processes, including autophagy and selective autophagy. An siRNA-based screen of the human kinome identified CK1α as one of ten kinases negatively regulating autophagy [26]. In addition, three independent studies have shown that CK1α phosphorylates the DEP- domain-containing mTOR-interacting protein (DEPTOR), promoting its proteasomal degradation and thereby facilitating full activation of mTOR signaling [27–29]. More recently, work from our team and others identified the selective autophagy receptor p62 as a substrate of CK1α. In the context of cGAS-STING pathway termination, CK1α thus contributes to the control of p62 abundance and promotes p62-mediated autophagic clearance of STING [30–32]. Recently, the immunomodulatory imide drug Lenalidomide and its derivatives, used in the clinic to treat multiple myeloma and myelodysplastic syndrome patients with deletion of chromosome 5q (del(5q)), were retrospectively characterized as small molecular glue degraders (MGDs) of CK1α. These MGDs directly connect CK1α to the E3 ubiquitin ligase cereblon (CRBN) for proteasomal degradation [33, 34].

Here, using multi-approach proteomics, we identify a lysosomal signature associated with CK1α. Through biochemical analyses, we show that CK1α depletion, either genetically or chemically, increases the overall abundance and maturation of proteins enriched in late endosome-lysosomes. This lysosomal remodeling is further associated with impaired tumorsphere formation capacity in GSCs. Using MGDs, we also uncover an additional role for CK1α associated with GPX4 inhibitor-induced ferroptosis, in both GSCs and THP1 cells, identifying CK1α as a regulator of ferroptotic sensitivity.

## Results

### Multi-proteomic profiling identifies a lysosomal signature associated with CK1α

We recently performed a comprehensive proteomic profiling in human monocytic THP1 cells depleted of CK1α (CK1α^sg^) using CRISPR-Cas9-mediated gene editing. Three datasets were generated: (i) a whole proteome analysis comparing control and CK1α-deficient cells, (ii) a proximity-based proteome (proximitome) using APEX2- tagged CK1α stably expressed in CK1α-deficient THP1 cells, and (iii) an interaction-based proteome (interactome) generated by immunoprecipitation of endogenous CK1α from WT and CK1α-KO THP1 cells. This multimodal approach led to the identification of CK1α as a key regulator of the cGAS-STING pathway [30] (**Fig. 1A**). Further analysis of the whole-proteome datasets revealed a strong enrichment of lysosome-associated proteins in CK1α^sg^-transduced cells across Gene Ontology (GO) Cellular Component (CC) and Molecular Function (MF) categories, as well as in the Kyoto Encyclopedia of Genes and Genomes (KEGG) pathway database (**Fig. 1B-C**). Consistently, lysosome-associated proteins were enriched in both the proximity and interaction networks of CK1α, suggesting that the kinase could be physically and functionally linked to lysosomal compartments (**Fig. 1D-E**). Overall, the joint analysis of these three complementary proteomic datasets points to a previously underestimated connection between CK1α and lysosomal biology.

**Figure 1:**
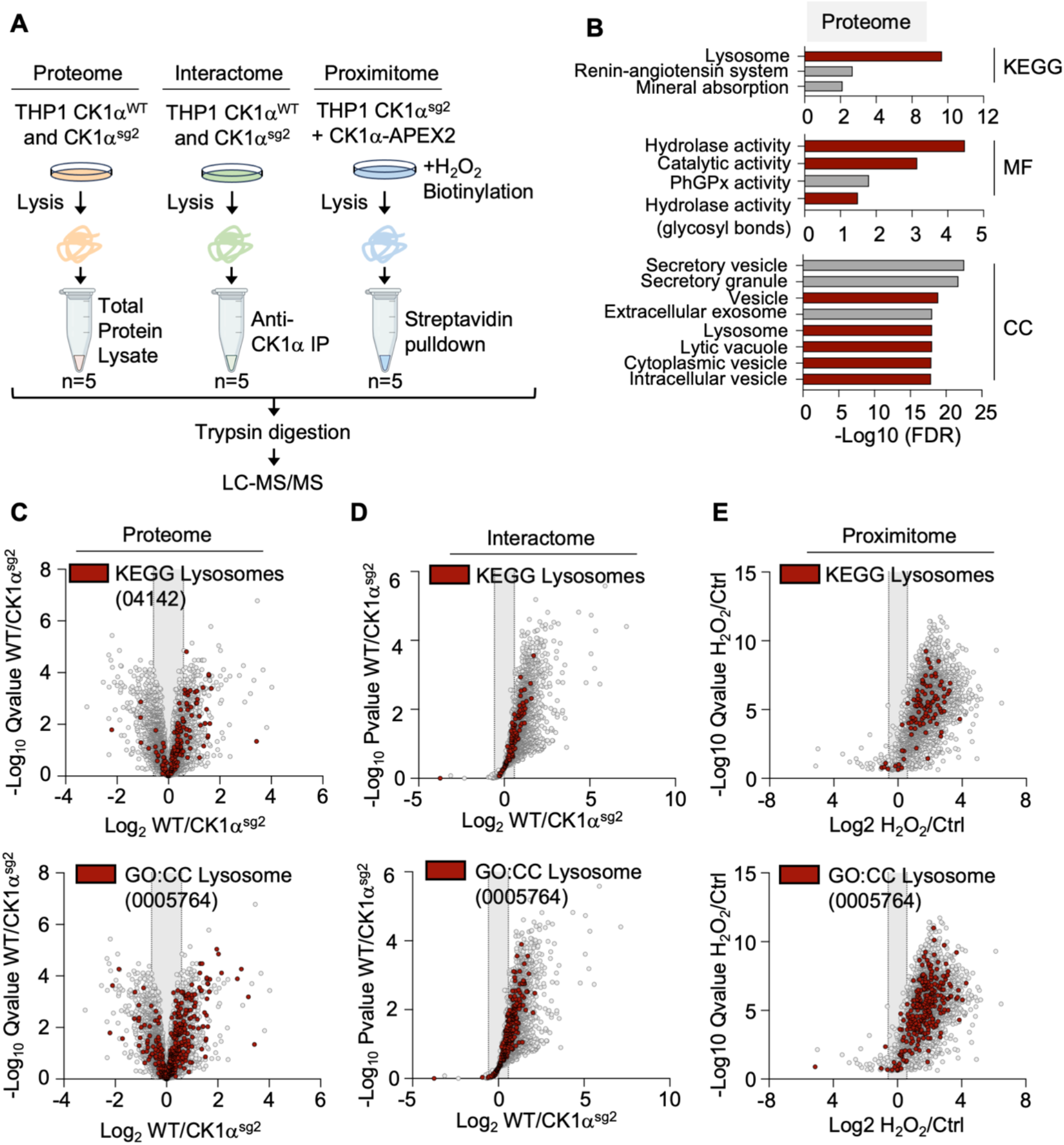
Convergent proteomic analyses identify a lysosomal signature associated with CK1α. (**A**) Schematic flow chart of the multi-omic approach performed in THP1 cells depleted of CK1α (CK1α ^sg^). (**B**) Functional enrichment analysis using Gene Ope-style/Group Statistics (g:GOSt) on g:Profiler of the top 100 upregulated proteins (qValue> 0.05) identified in the whole proteome dataset of THP1 CK1α ^sg^/ THP1 CK1α ^WT^ cells. (**C-E**) Gene Ontology analysis of lysosome-associated proteins from the KEGG lysosome pathway (KEGG:04142) or from the Cellular Compartment term lysosome (GO:CC 0005764) within (C) the THP1 CK1α^sg^ / CK1α^WT^ whole proteome, (D) the THP1 CK1α interactome, and (E) the THP1 APEX2-CK1α proximitome. Data are representative of five independent experiments.

### Genetic depletion of CK1α induces lysosomal remodeling

To investigate the impact of CK1α on lysosomal biology, we first assessed the levels of key lysosomal proteins, including the Lysosome-Associated Membrane Proteins 1 and 2 (LAMP1, LAMP2) and the lysosomal lumen aspartyl protease Cathepsin D (CTSD) by western blotting (**Fig. 2A**). During their maturation process, LAMP1 and LAMP2 undergo extensive N- and O-linked glycosylation, whilst CTSD is synthesized as a precursor that undergoes sequential proteolytic processing to generate its mature form [35]. In CK1α-depleted THP1 cells, both the glycosylated and unglycosylated forms of LAMP1 and LAMP2 proteins, as well as the mature form of CTSD, were more abundant. Accordingly, confocal microscopy revealed increased LAMP2 staining intensity in the absence of CK1α (**Fig. 2B-C**). This suggests an overall increase in both the abundance of proteins enriched in late endosome-lysosomes and their maturation in lysosomes. To further interrogate lysosomal function, we next employed LysoTracker, a fluorescent probe that selectively stains acidic intracellular compartments *in cellulo.* A two-fold increase in the mean fluorescence intensity (MFI) was robustly observed in cells lacking CK1α (**Fig. 2D**).

**Figure 2:**
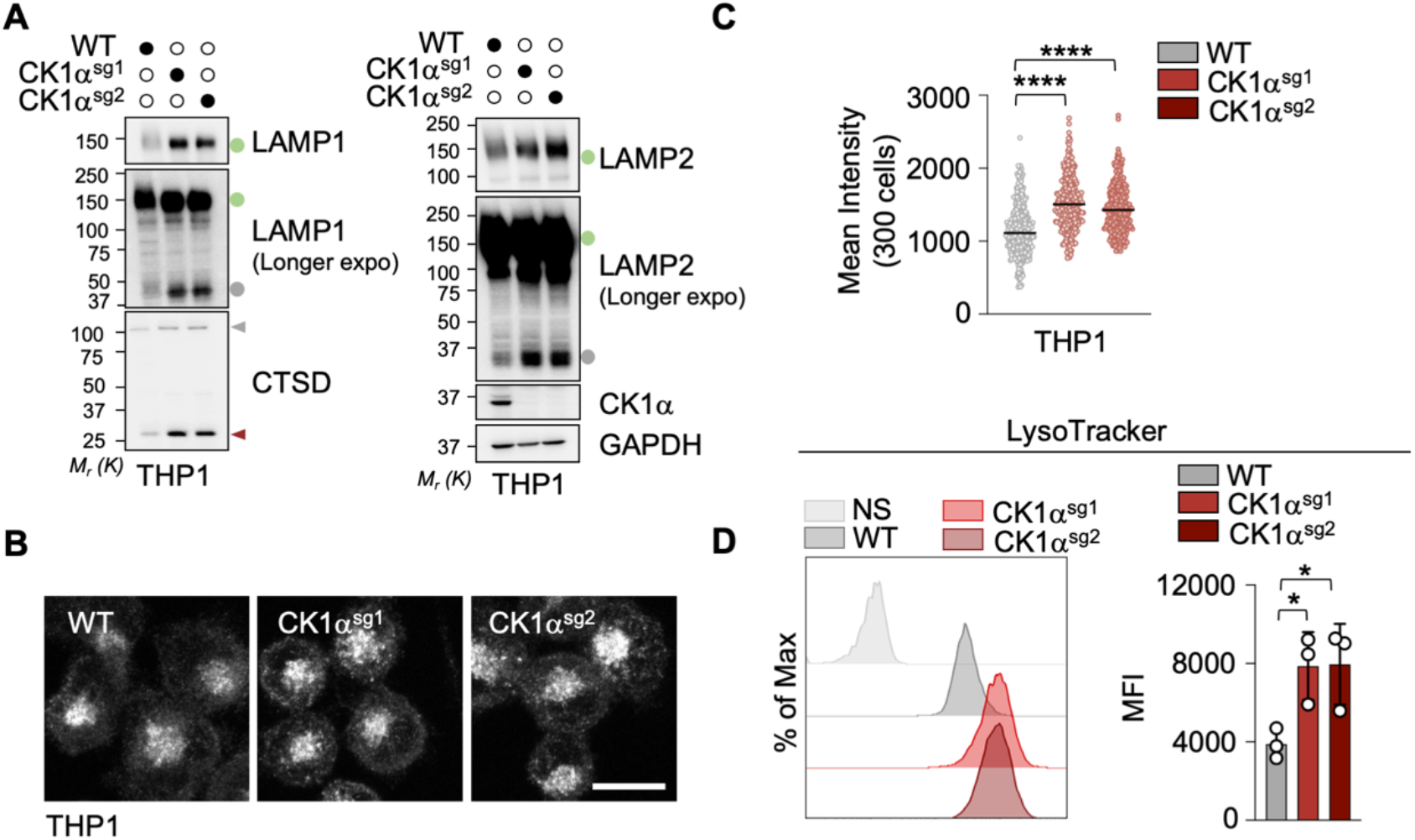
CK1*a* governs lysosomal fitness in THP1 cells. (**A**) Lysosomal protein abundance and post-translational modifications were assessed by immunoblotting in CK1α sgRNA-transduced THP1 cells. (**B-C**) Lysosomes were visualized by immunofluorescence using an anti-LAMP2 antibody in CK1α sgRNA-transduced THP1 cells. (B) Representative confocal microscopy images (Scale bar, 10µm). (C) Quantification of mean LAMP2 fluorescence intensity per cell. (**D**) Acidic compartments, including lysosomes, were stained with LysoTracker and quantified by flow cytometry in THP1 cells transduced with CK1α-targeting sgRNA or control sgRNA. (C-D) Statistical analyses were performed using ordinary one-way ANOVA followed by Dunnett’s multiple comparisons test (*p<0.05, **** p<0.0001). Data are representative of at least three independent experiments.

Given the emerging role of lysosomes as critical determinants of glioblastoma cell fate beyond metabolism [10, 36], we next investigated whether CK1α could be operating in the glioblastoma stem-like cell population (GSC). We therefore hypothesized that interfering with CK1α might disrupt their lysosomal homeostasis and, subsequently, compromise their fitness. We observed an increase in glycosylated forms of LAMP1 and LAMP2, together with enhanced expression of mature CTSD upon siRNA- mediated depletion of CK1α in two patient-derived GSCs, namely mesenchymal and classical GSC6 and GSC9 from 68-year-old male and female patients, respectively (**Fig. 3A**). Moreover, both cell types displayed increased LysoTracker staining following CK1α silencing, consistent with changes in the acidic endolysosomal compartment (**Fig. 3B**). Of note, the capacity of CK1α-silenced GSCs to form tumorspheres was decreased. In addition, CellTiter Glo assays revealed a reduction in metabolic viability without CK1α (**Fig. 3C-E**). Overall, these results suggest that CK1α regulates the lysosomal compartment in both THP1 cells and GSCs.

**Figure 3:**
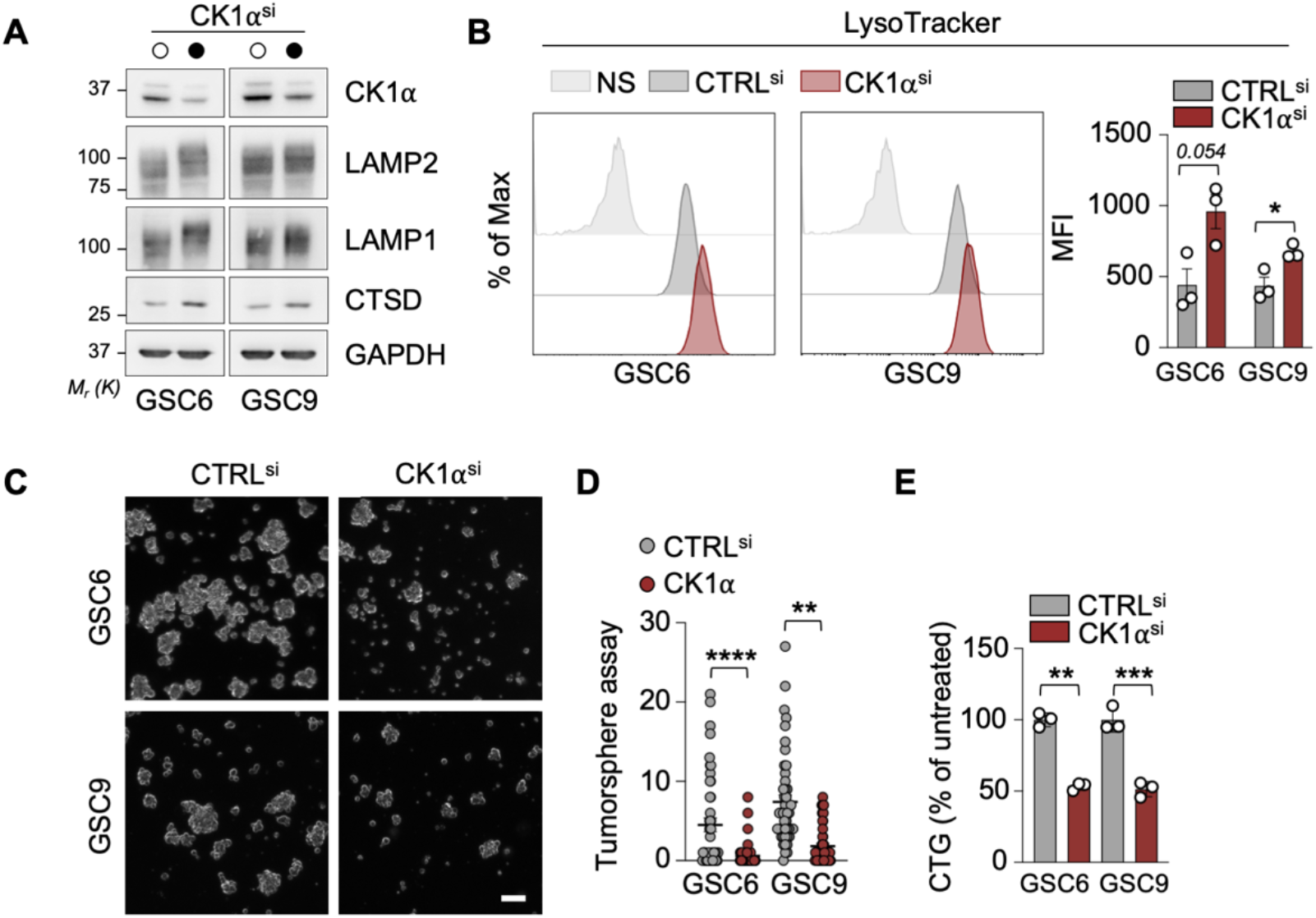
Targeting CK1*a* impairs patient Glioblastoma Stem-like Cells lysosomal fitness, tumorsphere formation, and viability. (**A-E**) GSC6 and GSC9 cells were transfected with siRNA targeting CK1α or a control nonspecific siRNA for 72h. (**A**) Lysosomal protein abundance and post-translational modification were assessed by immunoblotting. (**B**) Acidic compartments, including lysosomes, were stained with LysoTracker and quantified by flow cytometry. (**C**) Representative brightfield microscopy images. Scale bars, 100 µm. (**D**) Single-blinded quantification of tumorosphere numbers per field of view (15 fov per condition). (**E**) Cell viability was assessed using the ATP-based CellTiter-Glo assay. Data are presented as mean ± s.e.m. (n=3). (B-E) Statistical analyses were performed using ordinary one-way ANOVA followed by Dunnett’s multiple comparisons test (*p<0.05, **p<0.01, ***p<0.001, **** p<0.0001). Data are representative of at least three independent experiments.

### Pharmacological elimination of CK1α using molecular glue degraders induces lysosomal remodeling

To next evaluate whether acute depletion of CK1α protein recapitulates the lysosomal phenotypes observed with its silencing, we used two recently characterized next- generation MGDs, namely DEG-77 and SJ3149. They are both derived from Lenalidomide and use the E3 ubiquitin ligase CRBN to target CK1α for proteasomal degradation [30, 33, 34] (**Fig. 4A**). Treatment with either MGD resulted in efficient degradation of CK1α in both GSCs, similar to the effect observed with SJ3149 in THP1 cells (**Fig. 4B-C**). This is accompanied by an altered pattern of glycosylated LAMP1 and LAMP2 (**Fig. 4B-C**). Likewise, confocal analysis revealed a marked increase in LAMP2 staining in both cell types (**Fig. 4D-F**). Further supporting lysosomal remodeling, Lysotracker-positive acidic compartments were enhanced in MGD-treated THP1, GSCs, and the human glioblastoma cell lines U87 and U118 (**Fig. 4G-I**, **S1A- B**). Similar to CK1α silencing, the tumorsphere formation capacity of GSCs was impaired, although metabolic viability was only modestly reduced (**Fig. 4J-K**). Of note, MGD-induced lysosomal remodeling did not increase GSC sensitivity to lysosomal membrane permeabilization triggered by L-leucyl-L-leucine methyl ester (LLOMe) (**Fig. 5A-B**). Hence, treatment with CK1α MGDs phenocopies the lysosomal perturbation and self-renewal disruption observed following genetic or siRNA-mediated deletion, without impacting sensitivity to LLOMe-mediated cell death, further supporting a role for CK1α in maintaining lysosomal homeostasis.

**Figure 4:**
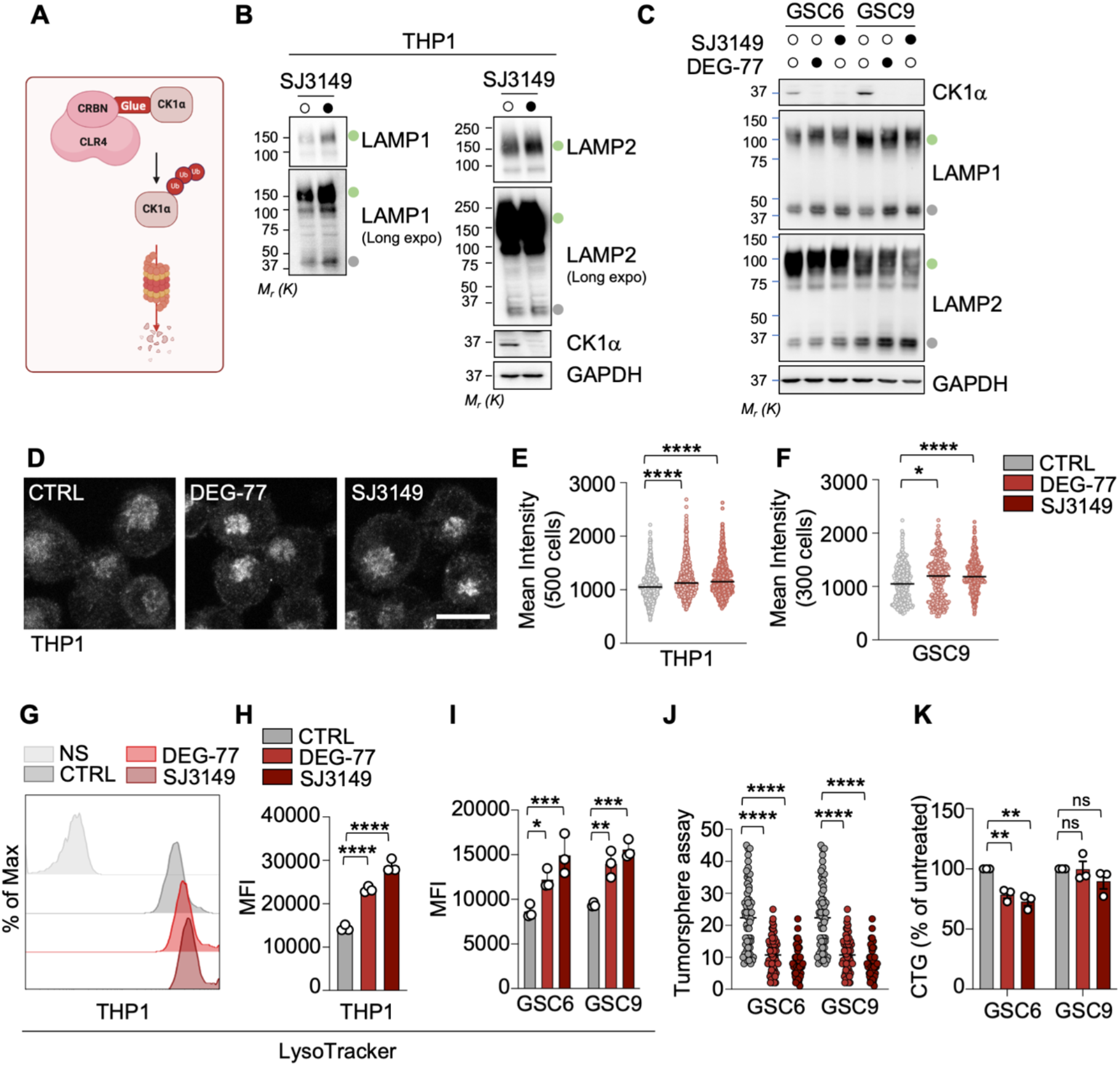
Molecular glue degraders disrupt lysosomal fitness in patient-derived glioblastoma stem-like cells. (A) Schematic representation of the mode of action of MGDs. (**B-C**) Lysosomal protein abundance and post-translational modifications were assessed by immunoblotting in (B) THP1 treated for 48h with 1µM SJ3149, and (C) the indicated GSCs treated for 72h with DEG-77 or SJ3149 (both at 1µM). Green circles indicate the O- and N- glycosylates form of the LAMPs proteins. Grey circles point to their unglycosylated forms. (**D-F**) Lysosomes were visualized by confocal microscopy by staining for LAMP2 by immunofluorescence in THP1 or GSC cells treated for 72h with 1µM DEG- 77 or SJ3149. (D) Representative confocal microscopy images for the THP1 cells dataset (Scale bar, 10µm). (**E-F**) Quantification of mean LAMP2 fluorescence intensity per cell in (E) THP1 cells or (F) GSC9 cells. (**G-I**) Acidic compartments, including lysosomes, were stained with LysoTracker and quantified by flow cytometry in THP1 (G-H) or in GSC6 and GSC9 (I) cells treated as in (E-F). (**J**) Single-blinded quantification of tumorosphere numbers per field of view (15 fov per conditions) from GSC6 and GSC9 cells treated with SJ3149 or DEG-77 (1µM, 72h). (**K**) Cell viability was assessed using the ATP-based CellTiter-Glo assay in GSCs treated as in (J). Data are presented as mean ± s.e.m. (n=3). (E, F, H) Statistical analyses were performed using ordinary one-way ANOVA followed by Dunnett’s multiple comparisons test. (I, J, K) Statistical analyses were performed using ordinary two-way ANOVA followed by Tukey’s multiple comparisons test (*p<0.05, **p<0.01, ***p<0.001, **** p<0.0001). Data are representative of at least three independent experiments.

**Figure 5:**
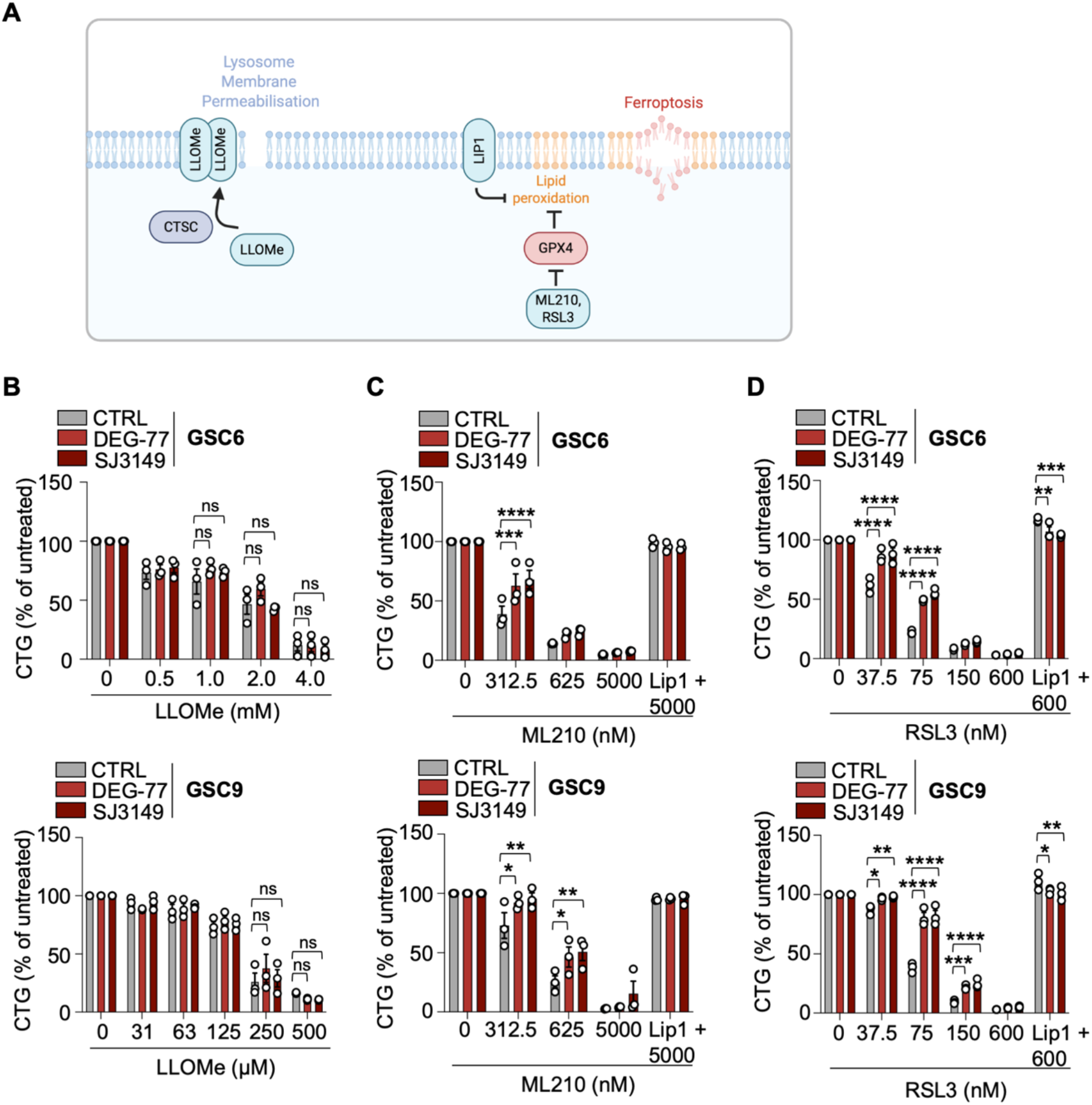
Molecular glue degraders protect both GSCs and THP1 cells from ferroptosis. (**A**) Schematic representation of the mechanisms of action of LLOMe in lysosomal membrane permeabilisation (LMP), and Liproxstatin-1 (Lip1), ML210 and RSL3 in ferroptosis. LLOMe is processed by cathepsin C (CTSC) within lysosomes, generating peptide oligomers that promote lysosomal membrane damage. ML210 and RSL3 induce ferroptosis by inhibiting glutathione peroxidase 4 (GPX4), thereby promoting iron-dependent lipid peroxidation. Conversely, Lip1 inhibits ferroptosis by inactivating lysosomal iron and preventing lipid peroxidation. (**B**) GSC6 and GSC9 cells were pretreated with the CK1α- targeting MGDs, DEG-77 and SJ3149 (1µM, 48h). Lysosome membrane permeabilization was induced with escalating concentrations of LLOMe for 16h. (**C**) GSC6 and GSC9 cells were pretreated with the CK1α- targeting MGDs, DEG-77 and SJ3149 (1µM, 72h). Ferroptosis was induced with escalating concentrations of the GPX4 inhibitor ML210 for 16h. When indicated, cells were pretreated for 2h with 1 µM of the ferroptosis inhibitor Liproxstatin-1 (Lip1). Cell viability was assessed using the ATP-based CellTiter-Glo assay. (**D**) CellTiter-Glo assays in GSC6 and GSC9 cells treated with another GPX4 inhibitor, RSL3, at the indicated concentration for 16h and co-treated, when indicated, with Lip1 (1µM). Cells were pretreated with DEG-77 and SJ3149 (1µM, 48h) before induction of cell death. Data are mean ± SEM of 3 biological repeats (*p<0.05, **p<0.01, ***p<0.001, **** p<0.0001; Ordinary two-way ANOVA followed by Dunnett’s multiple comparisons test).

### CK1α modulates the GSC ferroptotic sensitivity

Lysosomes play an important role in cellular iron homeostasis and have been implicated in the regulation of ferroptosis [18]. Interestingly, CK1α itself has recently been associated with ferroptosis. The CK1α E98V gain-of-function mutation associated with del(5q) myelodysplastic neoplasms was reported to deregulate iron handling and render bone marrow cells more sensitive to ferroptosis [37]. These observations, together with the lysosomal phenotype associated with CK1α, prompted us to investigate whether CK1α regulates ferroptotic sensitivity in GSCs. To this end, ferroptosis was triggered using two structurally distinct GPX4 inhibitors, RSL3 and ML210, which provoke iron-dependent membrane lipid peroxidation (**Fig. 5A**). Both compounds markedly reduced GSC metabolic survival at nanomolar concentrations, and this effect was rescued by the radical-trapping antioxidant Liproxstatin-1 (Lip1), confirming that they were undergoing ferroptotic death (**Fig. 5C-D**). It is noteworthy that GSCs underwent cell death by ferroptosis only when cultured in the presence of FBS, but not serum-free mitogen neurobasal medium (**Fig. S2A**). Strikingly, degradation of CK1α with either SJ3149 or DEG-77 significantly improved the survival of GSCs treated with RSL3 or ML210 (**Fig. 5C-D**). These findings were recapitulated in THP1 cells following genetic depletion of CK1α or treatment with MGDs (**Fig. S2B- E**). Thus, loss of CK1α protects GSCs and THP1 cells from GPX4 inhibitor-induced ferroptosis, identifying CK1α as a regulator of ferroptotic sensitivity. Altogether, these findings suggest that CK1α -dependent lysosomal modeling may fine-tune cellular susceptibility to ferroptosis.

## Discussion

CK1α has previously been implicated in the regulation of autophagy and lysosome-associated signaling [26–29]. Yet, how precisely CK1α affects the lysosomal compartment continues to be elucidated. Using multi-proteomics approaches, we now report that CK1α stands in proximity to lysosomal proteins and that its depletion is associated with broad changes in the lysosomal proteome. Both genetic ablation and acute MGD-mediated degradation of CK1α changed the maturation of lysosomal proteins, increased LysoTracker fluorescence, and impaired cell death by ferroptosis in two patient-derived GSCs. This effect was recapitulated in a monocytic cell line and an established glioblastoma cell line. Nevertheless, the molecular mechanisms underlying this lysosomal remodeling and the direct substrates of CK1α remain to be determined.

One potential mechanism may involve the master transcriptional regulator of lysosomal biogenesis and autophagy Transcription Factor EB (TFEB) [38]. Accordingly, TFEB was recently shown to bind the β-catenin destruction complex, which also involves CK1α, and regulate a subset of Wnt-dependent genes [39]. Alternatively, CK1α may act through the RNA-binding protein QKI, which maintains GSC stemness by limiting endolysosomal abundance and thereby reducing the degradation of self-renewal-promoting receptors at the plasma membrane [11]. Another possibility involves the paracaspase MALT1. Others and we previously showed that CK1α enables the activation of MALT1 and subsequent proteolysis of its dedicated substrates in immune cells [23, 24]. Inhibiting the protease activity of MALT1 was reported to increase the abundance of the endo-lysosome compartment, disrupt intracellular cholesterol homeostasis, impair autophagy, and promote lysosome-mediated cell death [13, 14]. Whether CK1α functions in coordination with MALT1 will deserve additional work.

We found that the GPX4 inhibitors RSL3 and ML210 induced potent ferroptosis in GSCs. This effect was observed only in the presence of serum, suggesting that GSC sensitivity to ferroptosis is strictly dependent on extrinsic factors, such as iron and cystine [40, 41]. This reiterates the potential contribution of extracellular signals, and by extension, the tumor microenvironment (TME) in determining the ferroptosis sensitivity. The GBM microenvironment is highly complex, comprising cellular (immune cells, blood vessels, astrocytes) and non-cellular (extracellular matrix and cues) components, which collectively create distinct tumor habitats. GSCs actively interact with and shape these niches, while their stem-like properties are themselves modulated by signals from the surrounding microenvironment [42]. Accordingly, our findings raise the possibility that GSC susceptibility to ferroptosis may vary across specific metabolic or microenvironmental niches, associated with cell states [43]. This is in line with the recent description of ferroptosis selectivity in the course of melanoma cell transition [44].

Ferroptosis primary initiates in the lysosomes and requires a functional lysosomal compartment, dynamically linked to organelles [18, 44, 45]. Consistent with this, treatment with Bafilomycin A1 or hydroxychloroquine, which increases the lysosomal pH, reduces ferroptosis [17, 18]. Our findings suggest that this could occur without increasing pH. CK1α degradation with MGDs remodels the lysosomal proteome and reduces the sensitivity of GSCs to ferroptosis inducers. This observation is consistent with a role of lysosomes in influencing ferroptosis susceptibility through broader mechanisms. Lysosomes are also the site of ferritinophagy, contribute to GPX4 turnover through chaperone-mediated autophagy, and regulate cellular glutathione and lipid metabolism [15]. The latter is particularly relevant, as the cellular threshold for ferroptosis induction can be determined by the relative abundance of distinct lipid species [46]. Lysosomal signaling may therefore indirectly influence ferroptotic susceptibility through metabolic pathways. For example, mTORC1 activation at the lysosome promotes SREBP1-dependent synthesis of monounsaturated fatty acids (MUFAs), which can protect cells from ferroptosis. Determining how CK1α-dependent lysosomal remodeling affects these metabolic pathways will therefore be important to understand how CK1α influences ferroptotic sensitivity in GSCs. Finally, small-molecule CK1α activators, such as pyrvinium and SSTC3, have been reported to enhance CK1α stability and activity through direct binding to the kinase [47]. It would therefore be interesting to determine whether pharmacological activation of CK1α sensitizes GSCs to ferroptosis.

Our results show that next-generation CK1α-targeted MGDs efficiently deplete CK1α in patient-derived GSC models. Targeted protein degradation (TPD) has emerged as a rapidly expanding therapeutic strategy, offering new opportunities to selectively eliminate previously challenging protein targets. Highly potent and selective CK1α MGDs have proven efficacy in preclinical models of Acute Myeloid Leukemia (AML) [33, 34]. Moreover, CK1α degradation can be further enhanced by the small molecule SB-405483, which binds to a previously unknown allosteric pocket of CRBN and increases its capacity to degrade CK1α and other neosubstrates [48]. An important challenge will be the potential of MGDs to target proteins in the central nervous system. In that view, a potent and selective, brain-penetrant MGD targeting ALK was recently reported [49]. In mouse intracranial xenograft models of non-small cell lung cancer (NSCLC), this MGD induced potent tumor regression, providing proof of concept that molecular glue degraders can achieve therapeutically relevant activity against targets within the brain [49]. Consistent with this idea, Lenalidomide was shown to penetrate cerebrospinal fluid in patients with refractory central nervous system (CNS) lymphomas in a pilot study [50]. Together, these advances support the feasibility of exploring CK1α-targeted protein degradation as a strategy for glioblastoma.

## Material and Methods

### Cell culture conditions

THP1 cells were purchased from the American Type Culture Collection and were cultured in RPMI 1640 supplemented with high FBS (20%), Glutamax, non-essential amino acids, Hepes, sodium pyruvate, penicillin, streptomycin, and normocin (Invivogen). The THP1 transduced with CK1α-sgRNA cell line and CK1α-deficient THP1 cell line stably expressing APEX2-tagged CK1α were previously described in [30]. The two patient-derived GSC cell lines annotated as GSC6 (mesenchymal, 68- year-old male) and GSC9 (classical, 68-year-old female) were isolated and characterized, as described previously [51]. These cells were routinely cultured in sphere-forming conditions using the NS34 medium: DMEM-F12 containing Glutamax, penicillin, streptomycin, amphotericin B, and N2, G5, and B27 supplements. For the ferroptosis experiments, GSCs were grown in NS34 media without N2, G5, and B27 supplements and with 10% FBS added. U87 and U118 cells were purchased from the American Type Culture Collection and were cultured in DMEM supplemented with FBS (10%), non-essential amino acids, penicillin and streptomycin. All reagents mentioned above were from Life Technologies, unless specified. All cells were maintained at 37°C in a humidified incubator with 5% CO_2_ whilst being routinely tested and found to be negative for mycoplasma contamination.

### Multi-omic analysis

The three complementary proteomic datasets, which include a whole proteome analysis comparing control and CK1α-deficient cells, a proximity-based proteome using APEX2-tagged CK1α stably expressed in CK1α-deficient THP1 cells, and an interaction-based proteome generated by immunoprecipitation of endogenous CK1α from WT and CK1α-KO THP1 cells, were previously published in [30]. Data can be found at https://www.ebi.ac.uk/pride/archive/projects/PXD063855 [52].

### siRNA transfection

Silencing by siRNA in GSCs was performed over 72h by lipofection using RNAiMax Lipofectamine (Life Technologies). Following manufacturer instructions, 0.8x10^6^ GSCs were transfected with 20 nM of the following siRNA duplexes: stealth non-silencing control (low-GC 12935111; Life Technologies) and CK1α 5’- GGCAAGGGCUAAAGGCUGCAACAAA-3’.

### Immunoblotting

Cells were collected and lysed in RIPA buffer (25 mM Tris HCl pH 7.4, 150 mM NaCl, 0.1% SDS, 0.5% Na-Deoxycholate, 1% NP-40, 1 mM EDTA) supplemented with 1x Halt Protease Inhibitor cocktail (ThermoFisher Scientific). Cell lysates were clarified by centrifugation at 10,000*g* to remove insoluble material. Protein samples were subsequently diluted into 2x Laemmli buffer (Life Technologies) containing either 10% 2-β-Mercaptoethanol or 100 mM Dithiothreitol and denatured at 95°C for 3 min. The samples were resolved onto 3-15% Tris-Acetate gels using the NuPAGE system (Invitrogen) and transferred onto nitrocellulose membranes (GE Healthcare). Membranes were then immunoblotted with the following primary antibodies: CTSD (610801, BD Biosciences), CK1α (ab223144, Abcam), GAPDH (SC-32233, Santa Cruz), LAMP1 (611042, BD Biosciences), and LAMP2 (SC-18822, Santa Cruz).

### Tumorsphere formation

Tumorsphere formation assays were conducted as described in [51]. GSCs (30,000 cells) were seeded in triplicate in 24-well plates with 500 µl media. Cells were dissociated manually daily over 3 days. The number of tumorspheres per field of view was assessed on day 5 by manual, single-blind count. In total, five randomly selected fields of view were quantified per condition.

### Cell viability assay

Metabolic cell viability was assessed using the CellTiter Glo Luminescent assay (Promega) following the manufacturer’s recommendation. GSCs and THP1 (5,000- 10,000 cells) were seeded in 96-well plates in a final volume of 100 µl. For the ferroptosis experiments, GSCs were cultured in DMEM-F12 containing Glutamax, penicillin, streptomycin, amphotericin B, and 10% FBS. The cells were pre-treated with 1 µM of MGDs targeting CK1α, DEG-77 (HY-157334, MedChemExpress) and SJ3149 (HY-160444, MedChemExpress), for 72h then co-treated for 16h with the ferroptosis agonists, ML210 (S0788, Selleckchem) and RSL3 (NB-64-24158, NeoBiotech) with or without 1 µM of the ferroptosis inhibitor Liproxstatin-1 (NB-64-13035, NeoBiotech).

### Flow cytometry

Lysosomal compartments were stained with 100 nM LysoTracker Green DND-26 (ThermoFisher) for 30 min at 37°C. Following a PBS wash, cells were analyzed by flow cytometry using a FACS Canto II (BD Biosciences; Cytocell Facility, SFR François Bonamy, France) or Cytoflex LX (Beckman Coulter, SFR ICAT, France) (for experiments with U87 and U118 cells). Data were processed using FlowJo V10 software.

### Immunofluorescence

THP1 cells were treated with 0.5 µM Phorbol 12-myristate 13-acetate (PMA) for 3 h to promote their adherence prior to immunofluorescence. Following two large PBS washes, 1x10^6^ cells were seeded onto 12 mm diameter coverslips fixed with 4% paraformaldehyde, then permeabilized with 0.2% Triton-X100. The coverslips were further incubated with 3% Bovine serum albumin for 20 min, stained with a LAMP2 antibody (SC-18822, Santa Cruz) for 1h, with secondary antibodies (A21121, Invitrogen), labelled with DAPI (1 µg/ml) for 5 min, and mounted onto glass slides using the prolong gold antifade mounting reagent (Invitrogen).

For GSCs, the coverslips were first incubated with 400 µl of 100 µg/ml poly-lysine for 30 min, then rinsed twice with PBS. GSCs (1x10^6^) were allowed to bind electrostatically to the coated coverslips for 15 min, followed by fixation, permeabilization, staining, and mounting onto glass slides, as per the protocol used for THP1 cells. Images were acquired by confocal microscopy (Nikon A1rHD LFOV) with a 60x oil-immersion objective at the Nikon Excellence Centre (Micropicell, UMS Biocore, Nantes, France), using 0.5 μm Z-intervals. All images were analyzed using the ImageJ software.

## Statements and Declarations

### Author Contributions

All authors contributed to the study conception and design. Conceptualization: JJ, MLG, JG, NB; Methodology: JJ, MLG, KT, FG; Investigation: JJ, MLG, AL, IV, TS, LM, RM, GAG, MK, NB; Data curation: FG; Formal analysis: JJ, MLG, KT, FG; Visualization: JJ; Supervision: GB, JG, NB; Project administration: GB, JG, NB; Funding acquisition: GB, JG, NB. All authors read and approved the final manuscript.

### Data Availability

The raw data and extracted proteomics data have been deposited in the ProteomeXchange Consortium via the PRIDE repository with the following dataset identifiers: PXD066072, PXD063855, and PXD066276, respectively.

### Ethics approval

This study was reviewed and approved by the institutional review board of Laennec Hospital, Nantes, France, and performed in accordance with the Helsinki protocol.

### Consent to participate

Informed consent was obtained from all patients before sample collection for diagnostic purposes.

## Acknowledgments

We thank all members of the SOAP laboratory for their help and insightful discussions, especially Célia Jaunasse for the expertise in bioinformatic analysis; Raphaël Rodriguez (Curie Institute) for helpful suggestions; the Cytocell and MicroPicell facilities (SFR Santé François Bonamy, Nantes, France) for the expertise.

## References

1. Louis DN, Perry A, Wesseling P, et al (2021) The 2021 WHO Classification of Tumors of the Central Nervous System: a summary. Neuro Oncol 23:1231–1251. 10.1093/neuonc/noab106

2. Price M, Ballard CAP, Benedetti JR, et al (2025) CBTRUS Statistical Report: Primary Brain and Other Central Nervous System Tumors Diagnosed in the United States in 2018– 2022. Neuro Oncol 27:iv1–iv66. 10.1093/neuonc/noaf194

3. Galli R, Binda E, Orfanelli U, et al (2004) Isolation and Characterization of Tumorigenic, Stem-like Neural Precursors from Human Glioblastoma. Cancer Res 64:7011–7021. 10.1158/0008-5472.CAN-04-1364

4. Singh SK, Hawkins C, Clarke ID, et al (2004) Identification of human brain tumour initiating cells. Nature 432:396–401. 10.1038/nature03128

5. Patel AP, Tirosh I, Trombetta JJ, et al (2014) Single-cell RNA-seq highlights intratumoral heterogeneity in primary glioblastoma. Science 344:1396–1401. 10.1126/science.1254257

6. Volovetz J, Berezovsky AD, Alban T, et al (2020) Identifying conserved molecular targets required for cell migration of glioblastoma cancer stem cells. Cell Death Dis 11:152. 10.1038/s41419-020-2342-2

7. Bao S, Wu Q, Sathornsumetee S, et al (2006) Stem Cell–like Glioma Cells Promote Tumor Angiogenesis through Vascular Endothelial Growth Factor. Cancer Res 66:7843– 7848. 10.1158/0008-5472.CAN-06-1010

8. Liu G, Yuan X, Zeng Z, et al (2006) Analysis of gene expression and chemoresistance of CD133+ cancer stem cells in glioblastoma. Mol Cancer 5:67. 10.1186/1476-4598-5-67

9. Chen J, Li Y, Yu T-S, et al (2012) A restricted cell population propagates glioblastoma growth after chemotherapy. Nature 488:522–526. 10.1038/nature11287

10. Merlet L, Le Guyon M, Gavard J (2026) A lysosomal requiem for glioblastoma cells. Trends in Cancer 12:386–389. 10.1016/j.trecan.2026.01.012

11. Shingu T, Ho AL, Yuan L, et al (2017) Qki deficiency maintains stemness of glioma stem cells in suboptimal environment by downregulating endolysosomal degradation. Nat Genet 49:75–86. 10.1038/ng.3711

12. Le Joncour V, Filppu P, Hyvönen M, et al (2019) Vulnerability of invasive glioblastoma cells to lysosomal membrane destabilization. EMBO Mol Med 11:EMMM201809034. 10.15252/emmm.201809034

13. Jacobs KA, André-Grégoire G, Maghe C, et al (2020) Paracaspase MALT1 regulates glioma cell survival by controlling endo-lysosome homeostasis. The EMBO Journal 39:e102030. 10.15252/embj.2019102030

14. Maghe C, Trillet K, André-Grégoire G, et al (2024) The paracaspase MALT1 controls cholesterol homeostasis in glioblastoma stem-like cells through lysosome proteome shaping. Cell Reports 43:113631. 10.1016/j.celrep.2023.113631

15. Dixon SJ, Olzmann JA (2024) The cell biology of ferroptosis. Nat Rev Mol Cell Biol 1–19. 10.1038/s41580-024-00703-5

16. Hangauer MJ, Viswanathan VS, Ryan MJ, et al (2017) Drug-tolerant persister cancer cells are vulnerable to GPX4 inhibition. Nature 551:247–250. 10.1038/nature24297

17. Torii S, Shintoku R, Kubota C, et al (2016) An essential role for functional lysosomes in ferroptosis of cancer cells. Biochem J 473:769–777. 10.1042/BJ20150658

18. Cañeque T, Baron L, Müller S, et al (2025) Activation of lysosomal iron triggers ferroptosis in cancer. Nature 642:492–500. 10.1038/s41586-025-08974-4

19. Mai TT, Hamaï A, Hienzsch A, et al (2017) Salinomycin kills cancer stem cells by sequestering iron in lysosomes. Nature Chem 9:1025–1033. 10.1038/nchem.2778

20. von Krusenstiern AN, Robson RN, Qian N, et al (2023) Identification of essential sites of lipid peroxidation in ferroptosis. Nat Chem Biol 19:719–730. 10.1038/s41589-022-01249-3

21. Amit S, Hatzubai A, Birman Y, et al (2002) Axin-mediated CKI phosphorylation of β- catenin at Ser 45: a molecular switch for the Wnt pathway. Genes Dev 16:1066–1076. 10.1101/gad.230302

22. Liu C, Li Y, Semenov M, et al (2002) Control of β-Catenin Phosphorylation/Degradation by a Dual-Kinase Mechanism. Cell 108:837–847. 10.1016/S0092-8674(02)00685-2

23. Bidère N, Ngo VN, Lee J, et al (2009) Casein kinase 1α governs antigen-receptor- induced NF-κB activation and human lymphoma cell survival. Nature 458:92–96. 10.1038/nature07613

24. Gehring T, Erdmann T, Rahm M, et al (2019) MALT1 Phosphorylation Controls Activation of T Lymphocytes and Survival of ABC-DLBCL Tumor Cells. Cell Reports 29:873–888.e10. 10.1016/j.celrep.2019.09.040

25. Elyada E, Pribluda A, Goldstein RE, et al (2011) CKIα ablation highlights a critical role for p53 in invasiveness control. Nature 470:409–413. 10.1038/nature09673

26. Szyniarowski P, Corcelle-Termeau E, Farkas T, et al (2011) A comprehensive siRNA screen for kinases that suppress macroautophagy in optimal growth conditions. Autophagy 7:892–903. 10.4161/auto.7.8.15770

27. Duan S, Skaar JR, Kuchay S, et al (2011) mTOR Generates an Auto-Amplification Loop by Triggering the βTrCP- and CK1α-Dependent Degradation of DEPTOR. Molecular Cell 44:317–324. 10.1016/j.molcel.2011.09.005

28. Gao D, Inuzuka H, Tan M-KM, et al (2011) mTOR Drives Its Own Activation via SCFβTrCP- Dependent Degradation of the mTOR Inhibitor DEPTOR. Molecular Cell 44:290–303. 10.1016/j.molcel.2011.08.030

29. Zhao Y, Xiong X, Sun Y (2011) DEPTOR, an mTOR Inhibitor, Is a Physiological Substrate of SCFβTrCP E3 Ubiquitin Ligase and Regulates Survival and Autophagy. Molecular Cell 44:304–316. 10.1016/j.molcel.2011.08.029

30. Jardine J, Tarrillon M, André-Grégoire G, et al (2026) The kinase CK1α coordinates the initiation and termination of the cGAS-STING pathway. Cell Death Differ 1–13. 10.1038/s41418-026-01844-0

31. Pan M, Hu T, Lyu J, et al (2023) CSNK1A1/CK1α suppresses autoimmunity by restraining the CGAS-STING1 signaling. Autophagy 0:1–18. 10.1080/15548627.2023.2256135

32. Watanabe Y, Tsujimura A, Taguchi K, Tanaka M (2017) HSF1 stress response pathway regulates autophagy receptor SQSTM1/p62-associated proteostasis. Autophagy 13:133–148. 10.1080/15548627.2016.1248018

33. Park S-M, Miyamoto DK, Han GYQ, et al (2023) Dual IKZF2 and CK1α degrader targets acute myeloid leukemia cells. Cancer Cell 41:726–739.e11. 10.1016/j.ccell.2023.02.010

34. Minzel W, Venkatachalam A, Fink A, et al (2018) Small Molecules Co-targeting CKIα and the Transcriptional Kinases CDK7/9 Control AML in Preclinical Models. Cell 175:171–185.e25. 10.1016/j.cell.2018.07.045

35. Settembre C, Perera RM (2024) Lysosomes as coordinators of cellular catabolism, metabolic signalling and organ physiology. Nat Rev Mol Cell Biol 25:223–245. 10.1038/s41580-023-00676-x

36. Jain A, Zoncu R (2026) Lysosomes as hubs of metabolic sensing and cellular homeostasis. Molecular Cell 86:533–552. 10.1016/j.molcel.2026.01.011

37. Schmitz S, Pearce JE, Martinez-Høyer S, et al (2026) Csnk1a1 E98 mutations rewire signaling and metabolism in del(5q) myelodysplastic neoplasms. Leukemia. 10.1038/s41375-026-03122-x

38. Ballabio A, Bonifacino JS (2020) Lysosomes as dynamic regulators of cell and organismal homeostasis. Nat Rev Mol Cell Biol 21:101–118. 10.1038/s41580-019-0185-4

39. Kim S, Song G, Lee T, et al (2021) PARsylated transcription factor EB (TFEB) regulates the expression of a subset of Wnt target genes by forming a complex with β-catenin- TCF/LEF1. Cell Death Differ 28:2555–2570. 10.1038/s41418-021-00770-7

40. Rodriguez R, Schreiber SL, Conrad M (2022) Persister cancer cells: Iron addiction and vulnerability to ferroptosis. Molecular Cell 82:728–740. 10.1016/j.molcel.2021.12.001

41. Ahmad F, Rendina BP, Chen C, et al (2026) Systemic cyst(e)inase administration induces ferroptosis and synergizes with temozolomide in glioblastoma. iScience 29:114350. 10.1016/j.isci.2025.114350

42. Bikfalvi A, da Costa CA, Avril T, et al (2023) Challenges in glioblastoma research: focus on the tumor microenvironment. Trends in Cancer 9:9–27. 10.1016/j.trecan.2022.09.005

43. Banu MA, Dovas A, Argenziano MG, et al (2024) A cell state-specific metabolic vulnerability to GPX4-dependent ferroptosis in glioblastoma. EMBO J 43:4492–4521. 10.1038/s44318-024-00176-4

44. Rizzollo F, Escamilla-Ayala A, Fattorelli N, et al (2025) BDH2-driven lysosome-to- mitochondria iron transfer shapes ferroptosis vulnerability of the melanoma cell states. Nat Metab 7:1851–1870. 10.1038/s42255-025-01352-4

45. Sassano ML, Tyurina YY, Diokmetzidou A, et al (2025) Endoplasmic reticulum– mitochondria contacts are prime hotspots of phospholipid peroxidation driving ferroptosis. Nat Cell Biol 27:902–917. 10.1038/s41556-025-01668-z

46. Magtanong L, Ko P-J, To M, et al (2019) Exogenous Monounsaturated Fatty Acids Promote a Ferroptosis-Resistant Cell State. Cell Chemical Biology 26:420–432.e9. 10.1016/j.chembiol.2018.11.016

47. Li B, Orton D, Neitzel LR, et al (2017) Differential abundance of CK1α provides selectivity for pharmacological CK1α activators to target WNT-dependent tumors. Science Signaling 10:eaak9916. 10.1126/scisignal.aak9916

48. Dippon VN, Rizvi Z, Choudhry AE, et al (2026) Identification of an allosteric site on the E3 ligase adapter cereblon. Nature 651:482–490. 10.1038/s41586-025-09994-w

49. Conery AR, La DS, Alekseyenko AA, et al (2026) TRI-611, a selective, brain-penetrant molecular glue degrader of ALK. Nature. 10.1038/s41586-026-10998-3

50. Rubenstein JL, Geng H, Fraser EJ, et al (2018) Phase 1 investigation of lenalidomide/rituximab plus outcomes of lenalidomide maintenance in relapsed CNS lymphoma. Blood Adv 2:1595–1607. 10.1182/bloodadvances.2017014845

51. Harford-Wright E, Andre-Gregoire G, Jacobs KA, et al (2017) Pharmacological targeting of apelin impairs glioblastoma growth. Brain 140:2939–2954. 10.1093/brain/awx253

52. Perez-Riverol Y, Bandla C, Kundu DJ, et al (2025) The PRIDE database at 20 years: 2025 update. Nucleic Acids Research 53:D543–D553. 10.1093/nar/gkae1011

