## Supplementary figures for "Molecular glue degraders of CK1α remodel lysosomes and limit ferroptosis in glioblastoma stem-like cells"

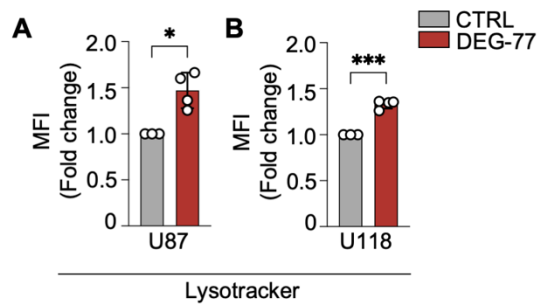

**Figure S1: Molecular glue degraders disrupt the lysosomal compartment in glioblastoma cell lines.**

(A-B) Acidic compartments, including lysosomes, were stained with LysoTracker and quantified by flow cytometry in U87 (A) or in U118 (B) cells treated with the CK1α-targeting MGD DEG-77 (2μM, 72h). Statistical analyses were performed using Welch's t-test (\*$p$<0.05, \*\*\*$p$<0.001). Data are representative of at least three independent experiments.

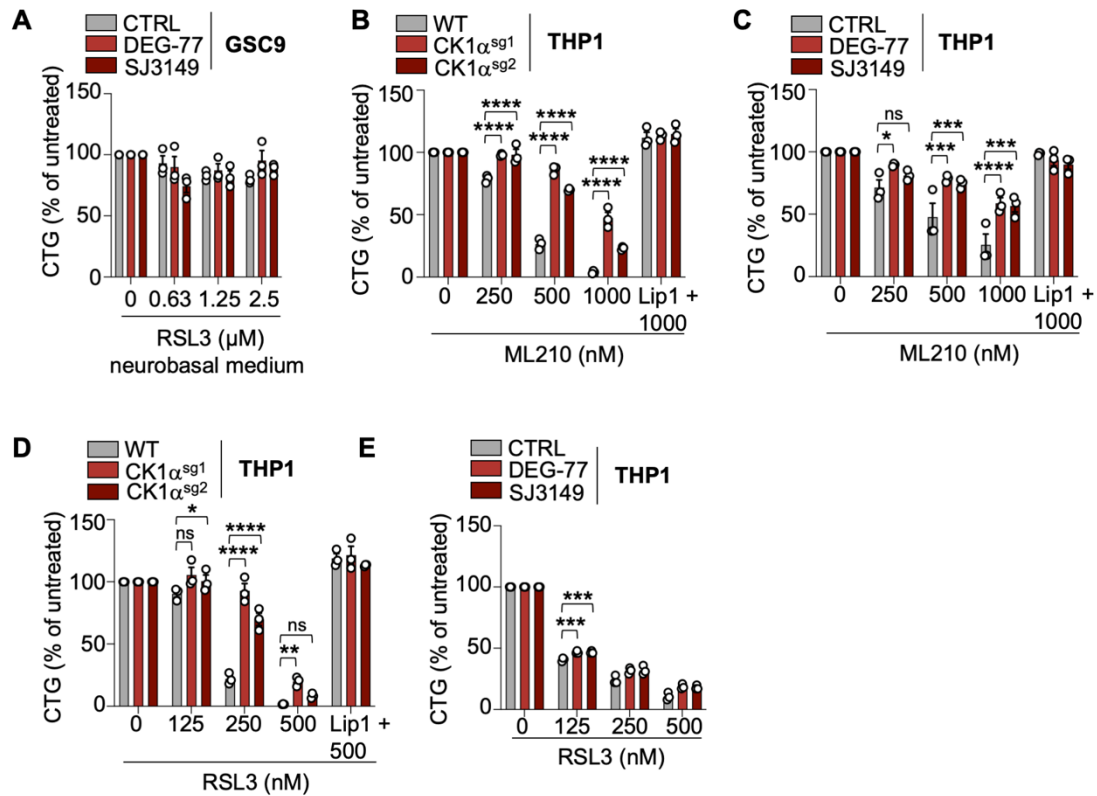

**Figure S2: Targeting CK1α protects THP1 cells from ferroptosis**

(A) GSC9 cultured in neurobasal medium were pretreated with the CK1α- targeting MGDs, DEG-77 and SJ3149 (1μM, 48h). Ferroptosis was induced with escalating concentrations of the GPX4 inhibitor RSL3 for 16h. (B-E) THP1 cells were transduced with CK1α sgRNA (B, D) or pretreated with the CK1α-targeting MGDs, DEG-77 and SJ3149 (1μM, 72h) (C, E). Ferroptosis was induced with escalating concentrations of the GPX4 inhibitor ML210 (B-C) or RSL3 (D-E) for 16h. When indicated, cells were pretreated for 2h with 1μM of the ferroptosis inhibitor Liproxstatin-1 (Lip1). Cell viability was assessed using the ATP-based CellTiter-Glo assay. Data are mean ± SEM of 3 biological repeats (\*p<0.05, \*\*p<0.01, \*\*\*p<0.001, \*\*\*\* p<0.0001; Ordinary two-way ANOVA followed by Dunnett's multiple comparisons test).
